# Adaptive transcranial ultrasound Doppler imaging of the brain

**DOI:** 10.1101/2025.05.27.656275

**Authors:** Rick Waasdorp, Eleonora Munoz Ibarra, Flora Nelissen, Baptiste Heiles, Valeria Gazzola, Guillaume Renaud, David Maresca

## Abstract

The development of fully noninvasive, transcranial functional ultrasound (fUS) would increase the translatability and clinical potential of this neuroimaging modality. Unfortunately, transcranial fUS is hindered by skull-induced aberrations which degrade power Doppler image quality and lower sensitivity. As a result, a majority of fUS imaging studies rely on craniotomies or acoustically transparent cranial windows. To advance fUS technology further, we present an adaptive aberration correction method based on ray-tracing through 4 tissue layers (transducer lens, gel & skin, bone and brain tissue). Our method segments these layers, and estimates ultrasound wave speeds in each layer iteratively. Once a velocity model of the imaging plane of interest is retrieved, ultrafast power Doppler imaging of the brain is performed using a ray-tracing beamformer that accounts for wave refraction. We tested our method in three adult rats, and estimated wave speeds for the skin/gel layer (1628 ± 7 m*/*s), skull bone (3247 ± 110 m*/*s), and brain tissue (1526 ± 55 m*/*s). After aberration correction, we measured an average adult rat skull thickness of 388 ± 41 μm in agreement with anatomical records. The largest improvements in Doppler imaging quality were observed in cortical brain layers adjacent to the skull, specifically lateral spatial resolution was improved by 32%. Our method consistently outperformed Doppler imaging based on traditional delay-and-sum (DAS) beamforming, which assumes a uniform sound speed.

## 1 Introduction

Functional ultrasound (fUS) has emerged as a valuable imaging modality for detecting cerebral blood volume variations associated with neural activity (1, 2). fUS stands out for its portability, sensitivity, and spatio-temporal resolution compared to functional magnetic resonance imaging (3). Clinically, fUS has been used at bedside to monitor neonate health (4) and intraoperatively in adults during tumor resection surgeries (5–7). fUS allows analysis of cerebral hemodynamic signals on a signal trial basis, making transient neural activity detectable (8). These capabilities led to decoding of movement intentions in nonhuman primates, which represented a fundamental step towards the development of ultrasound brain-computer interfaces (9). In spite of these advances, the skull remains a highly attenuating and aberrating barrier for ultrasound waves that prevents the development of fully non-invasive fUS and wider adoption of this technology.

The skull bone has a high density and speed of sound (typically 3000 m/s) compared to soft tissues (skin 1600 m/s, brain 1570 m/s) (10). As a result, transcranial ultrasound images are degraded by several physical effects including multiple scattering, mode conversion (11), absorption (12) and refraction (13) (Fig. 1). This currently restricts the use of fUS in adult humans to intraoperative procedures (7) or chronically implanted acoustic transparent windows (14). Transcranial fUS is feasible in mice and young rats due to the relatively thin skull (15, 16), but as animals age, and the skull develops (17), other solutions are needed. Most studies simply remove (1) or thin the skull (18) to prevent aberration, a procedure not compatible with clinical fUS or fUS based BMI. As an alternative to surgical procedures to the skull, the echogenicity of vascular signals can be enhanced using contrast agents (19, 20). However, the effect is transient due to gradual elimination of the agent, requiring continuous administration for sustained enhancement. Moreover, use of contrast agents is minimally invasive and only addresses ultrasound signal attenuation.

**Fig. 1.**
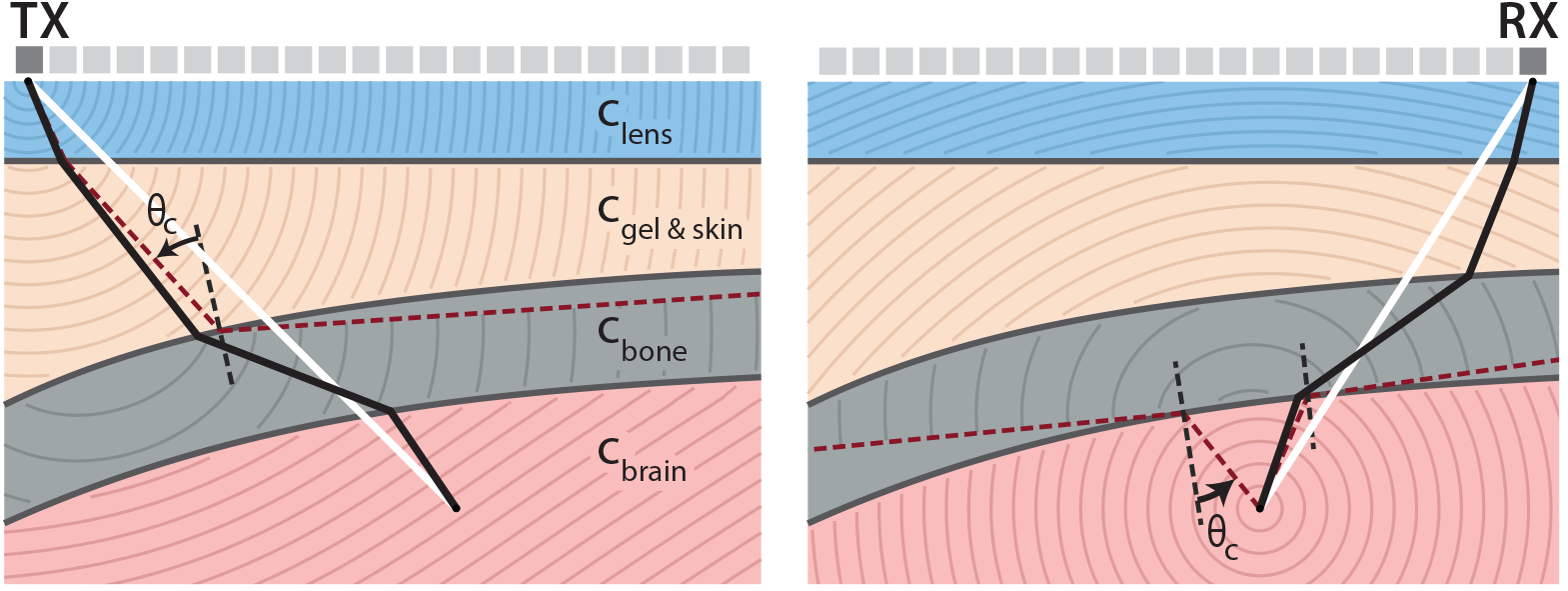
Wave refraction during transcranial ultrasound imaging. **(a)** Wave propagation in transmission and **(b)** reception, through 4 layers with distinct sound speeds: the transducer lens, acoustic coupling gel and skin tissue, the skull bone, and finally brain tissue. White lines show the wave path in conventional image reconstruction assuming homogeneous wave speed. Black lines show the refracted path through the medium, fulfilling Snell’s law. Iso-phase lines (wave front) in each layer are spaced at one wavelength *λ* in water. The dark red dashed lines indicate the paths subject to total internal reflection due to the critical angle *θ*_*c*_.

Alternative to addressing the attenuation problem, the loss of focus can be restored by aberration correction techniques (21). Most techniques rely on near field phase-screen modeling (22), where the aberrating layer is modeled as a thin phase screen located at the transducer surface. The phase aberration is corrected by finding the delays to apply to each transducer element, typically by maximizing the correlation between elements (23), maximizing spatial coherence (22), or maximizing speckle brightness (24). This method can be used in the presence of either bright reflectors such as microbubbles (25–27) or diffuse scatterers (28). Another class of aberration correction methods use ultrasound matrix imaging (29), which has been applied to transcranial ultrasound localization microscopy (ULM) (30). Finally, deep learning-based aberration correction (31, 32) has been developed but requires either contrast agents or has only been tested in weak aberrating conditions. However, all forementioned methods are limited in their ability to restore positional errors and the aspect ratio of the image.

Another group of methods uses a layered tissue velocity model to compute the refracted wave path and correct for phase aberration. This requires wave speeds for every tissue layer and an accurate delineation. The skull shape and wave speed can be obtained by ultrasound imaging (33–35), or by using other imaging modalities such as computed tomography (CT) (36–38). However, relying on other imaging modalities has the drawback of potentially introducing co-registration errors and the need for additional costly imaging systems. Once a complete sound speed map is obtained, the propagation of a wavefront through the medium can be computed using the Eikonal equation, and the aberration corrected time of flights are obtained. The Eikonal equation can be solved using the fast marching method (39). This has been used in simulation, phantoms (38, 40, 41), and *ex vivo* human skull (33, 42). Alternatively, the refracted wave path can be found using two point ray-tracing (43), allowing imaging of the bone cortex (34, 35), and *ex vivo* human skull imaging (42). The aberrated delays can be computed for every source-receiver pair, and finally be used in delay-and-sum (DAS) beamforming (44).

Here we present an adaptive transcranial ultrasound imaging method to estimate the rat skull geometry and wave speed, and corrects for wave refraction in transmission and reception using ray tracing. Compared to previously published methods, we do not require a priori information on the wave speeds or geometry of the different tissue layers, and use only ultrasound to identify aberrations induced by each layer on a per-subject basis. To evaluate the performance of this approach, we performed transcranial Doppler imaging in adult rat brains with and without aberration correction.

## 2 Methods

### 2.1 Animal procedures

Three adult male Sprague Dawley rats were imaged in this study (6-8 weeks old; 260-360 grams; Janvier Labs, France). Animals were socially housed in groups of 2-3 in a 12h reversed light-dark cycle and had ad libitum access to food and water. All experiments were approved under CCD license number AVD8010020209725 and performed at the Netherlands Institute for Neuroscience.

All ultrasound recordings were performed under anesthesia. Prior to imaging, the rats were anesthetized using 5% isoflurane in an induction chamber and then transferred to an inhalation mask with 3% isoflurane for depilating over the head. Next, the rats were placed on a heating pad set to 37 °C, and head-fixed in a stereotactic frame attached to an inhalation mask, delivering 1.5% isoflurane to maintain anesthesia. After positioning the rat in the imaging setup, anesthesia was transitioned from 1.5% isoflurane to medetomidine (0.1 mg/mL) via a single bolus injection. Three minutes later, isoflurane administration was discontinued, and deep anesthesia was confirmed by the absence of a toe pinch reflex. Subsequently, a continuous subcutaneous infusion of medetomidine (0.05 mg/mL) was initiated to maintain anesthesia throughout the imaging session. Centrifuged ultrasound gel (Ultrasonic 100, Parker Laboratories, Orange, NJ, USA) was applied to couple the scalp to the probe. The maximum duration of the experiment was 1.5 hours.

### 2.2 Ultrasound acquisitions

Ultrasound acquisitions were performed with a L6-24-D transducer (192 elements; 135 μm pitch; GE HealthCare, Chicago, IL, USA) connected to a programmable ultrasound scanner (Vantage 256 High frequency configuration, Verasonics, Kirk-land, WA, USA).

Our aberration correction method requires an accurate segmentation of the skull bone. To this end, we implemented a high resolution skull segmentation sequence that relies on synthetic aperture imaging with element-wise transmissions (45). To obtain the best axial resolution we transmitted short single-cycle sine-bursts centered at 13.9 MHz.

After bone image acquisition, we switched to plane wave imaging to acquire Ultrafast Power Doppler images. We transmitted 3 cycles sine-bursts centered at 13.9 MHz and 26 plane waves ranging from -5 to 5°, resulting in 700 Hz frame rate after coherent compounding (46). We set the duration of frame ensembles to 250 ms to capture a full cardiac cycle of the rat in each Doppler image.

To correct for refraction effects in the skull bone, the inner and outer skull interfaces must be clearly resolved in the B-mode. This requires to position the probe perpendicular to the skull surface to obtain specular echoes from both skull interfaces (see Fig. 2). We used live B-mode imaging to find maximal specular reflections from outer and inner skull interfaces. The probe was positioned 2 mm above the skull surface. Next, we acquired an ultrafast power Doppler image in the same plane using the plane wave sequence described previously.

**Fig. 2.**
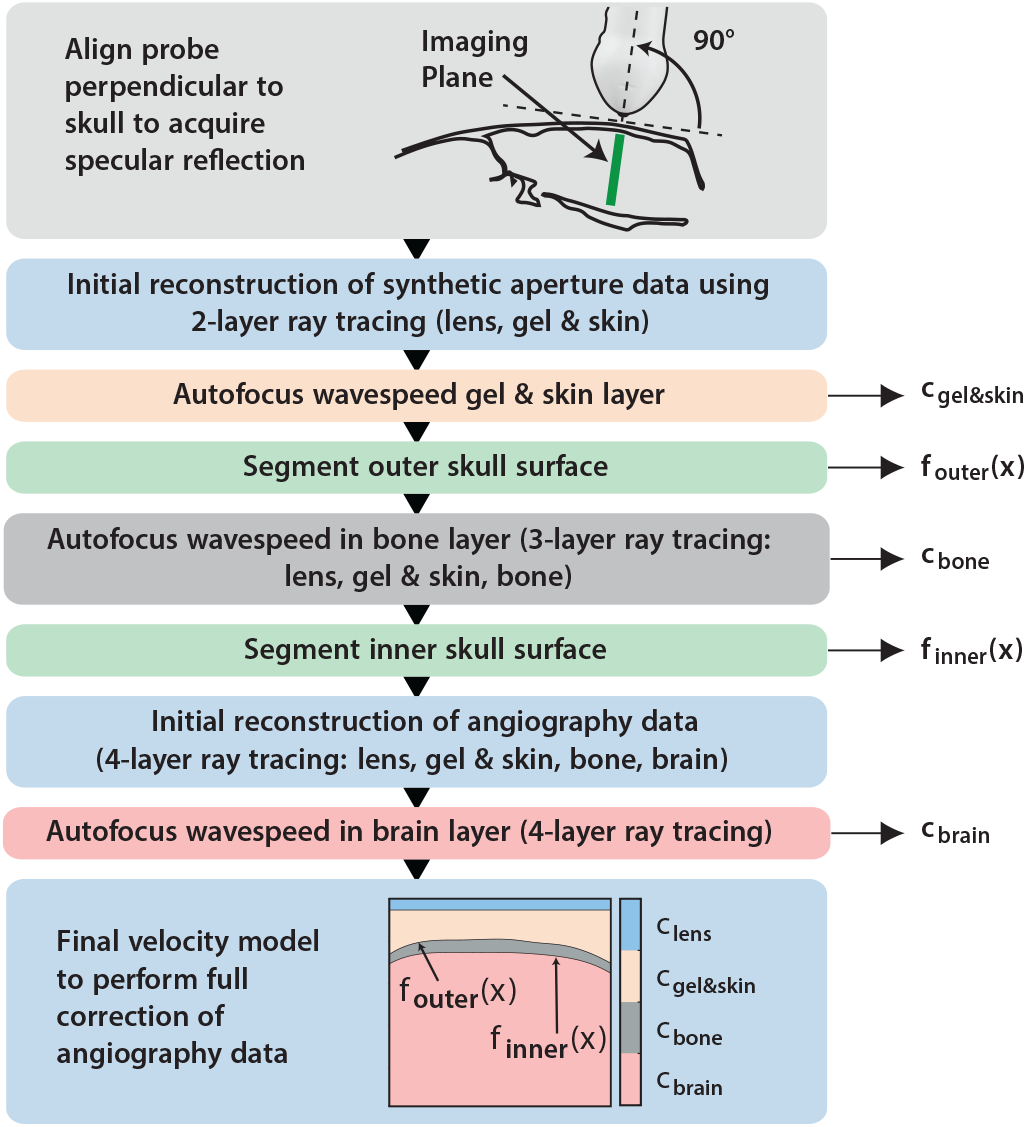
Graphical overview of the adaptive aberration correction method.

We used a real-time DAS (delay-and-sum) beamformer implemented in CUDA on a GPU (GeForce RTX 3090, NVIDIA, USA) for live display. After beamforming, power Doppler images were formed by separating blood signals from tissue signals using a singular value decomposition (SVD) spatio-temporal clutter filter (47). Since motion is minimal in head fixed anesthetized experiments, we used a fixed threshold and removed the first 20% of singular vectors (35 out of 175 frames). RF data from both skull segmentation and brain angiography sequences were saved and processed offline.

### 2.3 Delay-and-Sum Beamforming

The gold standard for ultrasound image reconstruction is DAS beamforming. For each pixel point, the time of flight of the ultrasound wave is calculated for each combination of transmitting and receiving elements. In conventional DAS, the medium is assumed to have a homogeneous sound speed *c*_0_ of 1540 m/s and straight paths are assumed between the source, pixel point and receiver (Fig. 1a). The image magnitude 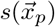 at pixel point 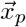 is given by,

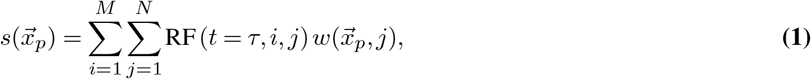

with *M* the number of transmissions, *N* the number of receiving elements, RF the recorded data, *τ* denotes the delay for pixel 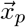, transmission *i* and receiving element *j*, and 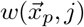 a weighting to account for element directivity (48).

Ultrasound scanners usually emit short bursts of sine waves, consisting of a few periods of oscillation at the probe’s central frequency. To ensure accurate image reconstruction, it is critical to delay the ultrasound signals to the crest of the envelope of the received waves, i.e. the time to peak *t*_ttp_ of the envelope of the received waveform (49). Finally, transducers have a lens to focus the ultrasound beam in the elevation plane. Lenses typically have a lower sound speed than soft tissue, and the difference in sound speed has to be accounted for. An estimation for the time to peak and the lens correction term is often given by transducer manufacturers.

To evaluate the aberration correction approach, we reconstructed reference images using the conventional DAS beamforming method with a homogeneous sound speed of 1540 m*/*s, manufacturer reported values for time to peak and lens correction, and a standard binary element weighting using F number 2 (44).

### 2.4 Aberration Correction Method

In transcranial imaging, the homogeneous wave speed assumption fails. The skull bone has a significantly higher wave speed than the soft tissues and therefore time of flights are underestimated using conventional DAS and wave refraction is not accounted for (Fig. 1a). To obtain accurate time of flights, we used an adaptive aberration correction method that estimates the wave speed in four tissue layers and corrects the time of flight by retrieving the refracted wave path through ray tracing (Fig. 1b).

The method is divided in three reconstruction steps. In the first step, we use a 2-layer model to correct lens refraction, which allows to accurately estimate the wave speed in the gel/skin layer by autofocusing (49), and segmentation of the outer skull surface. In the second step we use a 3-layer model to estimate the wave speed of the skull bone and segment the inner skull surface. During the final step, we use a 4-layer model to estimate the wave speed in the brain tissue and perform the final aberration correction. A graphical overview of the method is shown in Fig. 2.

#### 2.4.1 Transducer parameters: acoustic lens and time-to-peak

A prerequisite to all aberration corrections reported in the rest of the manuscript is to characterize the lens and waveform parameters of the transducer. We transmitted a 1-cycle imaging pulse in air in a synthetic aperture sequence, and received the internal lens reflections. Then we found the time to peak, lens thickness and lens wave speed by maximizing the coherence of the internal lens reflections as described in (49).

#### 2.4.2 4-Layer Velocity Model

The first layer models the transducer lens, and allows accounting for refraction at the lens interface. The second layer models both the acoustic coupling gel and skin tissue. The heterogeneity within this layer is assumed to be small, since ultrasound gel used for coupling is matched with skin tissue in terms of acoustic impedance. Skin wave speed is reported to be 1600 − 1620 m*/*s, slightly higher than soft tissue (10, 50).

The third layer in our model represents the skull bone, characterized by its high density and a reported wave speed ranging from 2400 to 3500 m*/*s (51–53). The thickness and curvature of the skull vary considerably, with the supraoccipital bone at the back measuring between 0.6 and 1.2 mm, and the parietal bone in the middle ranging from 0.2 to 0.6 mm, mainly depending on body mass (54). Rat skulls consist of plates that grow together and are connected by sutures. The speed of sound in the skull increases with age due to changes in mineral content of the cortical bone. The sutures consist of collagen-rich connective tissues with sound speeds closer to soft tissue. The skull bone is not a uniform layer but rather a sandwich structure consisting of an outer and inner cortical layer with diploë in between. The diploë is a spongy trabecular bone layer and typically constitutes about one-fifth of the total skull thickness (55, 56). Given that the diploë layer is too thin to resolve using ultrasound imaging and accounts for a small thickness fraction, we simplify the skull as a single homogeneous layer in our ray-tracing model. The varying curvature and thickness is modeled by segmenting the outer and inner surfaces using piece wise cubic splines.

The final layer models the brain tissue. There is no literature reported value for the wave speed in rat brain tissue at high frequency and *in vivo* temperatures. In fresh *ex vivo* canine brain samples, the wave speed was reported to be between 1563 − 1570m*/*s (57, 58), measured at 37 °C and 5 MHz, slightly higher than the averaged soft tissue value of 1540 m*/*s. We expect the rat brain wave speed to be in same range.

#### 2.4.3 Adaptive Ray Tracing

To find the aberrated round trip time of flight, we use a ray tracing method adapted from Renaud *et al*. (34) that finds the refracted wave path in the 4-layer model. The estimated time of flight is used in conventional DAS beamforming to reconstruct the corrected image (44).

The refracted path is found by two-point ray tracing, a technique that is commonly used in seismic imaging (43). In this technique, Snell’s law of refraction is imposed on all but the last interface. The aberrated wave path is found by minimizing the time of flight between a source and pixel point, or receiver and pixel point, following Fermat’s principle. Specifically, for each source and pixel combination, we test different emerging ray angles form the source, traverse the layers by computing the refracted ray at each interface intersection, except for the last. The last interface intersection point is connected to the end point, to compute a time of flight for the tested angle. Finally, we find the path fulfilling Snell’s law by minimizing the time of flight using Brent’s algorithm (59). This process is repeated for all combinations of sources and pixel points and receivers and pixel points.

The algorithm gives access to the reception angle for each source-pixel-receiver combination, which is used to account for element directivity in the beamforming process (48). The algorithm is implemented in C++ and parallelized using OpenMP.

#### 2.4.4 Intersection of rays with interfaces

During the minimization, the speed of sound map is traversed by rays with different angles. This requires finding the intersection points of a ray with the lens and skull surfaces. The intersection point on the lens surface follows from the tested ray angle and the lens thickness. The intersection point of the refracted ray with the outer skull surface is found by a line-spline intersection algorithm.

To find the intersection point of the ray and the piece wise cubic spline, we need to find in which subdomain the intersection point lies. This is done by testing the ray against all spline segments with potential intersections.

For a ray described by *z* = *px* + *q*, and a cubic spline section *z* = *ax*^3^ + *bx*^2^ + *cx* + *d*, we find their intersection point by computing the roots of the equation,

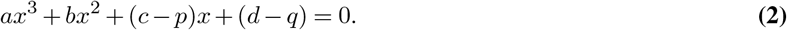

This cubic is reduced to a depressed cubic equation of the form,

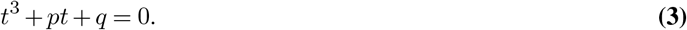

Finally, we apply Cardano’s formula to find only the real roots (60). We obtain 1 root if the discriminant is positive and 3 otherwise. Next we need to test if the found roots are in the domain of the current spline segment. The root is discarded if it is not in the domain, otherwise, we find the intersection point by evaluating the line equation at the root. If no valid real roots are found for a spline segment, the next segment is tested. If no valid roots are found for any segment, the ray does not intersect the spline.

#### 2.4.5 Wave speed estimation

We find the wave speed in each layer by autofocusing, in which we reconstruct a region of interest (ROI) of the aberrated image with different wave speeds, and select the wave speed that maximizes the image sharpness. Image sharpness is defined by the Brenner gradient,

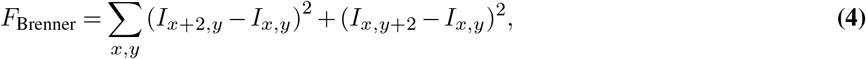

where *I*_*x,y*_ is the image intensity at location (*x, y*) (61).

The wave speed in the gel/skin layer is found by using the 2-layer model on the synthetic aperture data from the skull-segmentation sequence. After an initial reconstruction at 1600 m*/*s we define a region of interest (ROI) around the skull bone (Fig 4b) and autofocus.

**Fig. 3.**
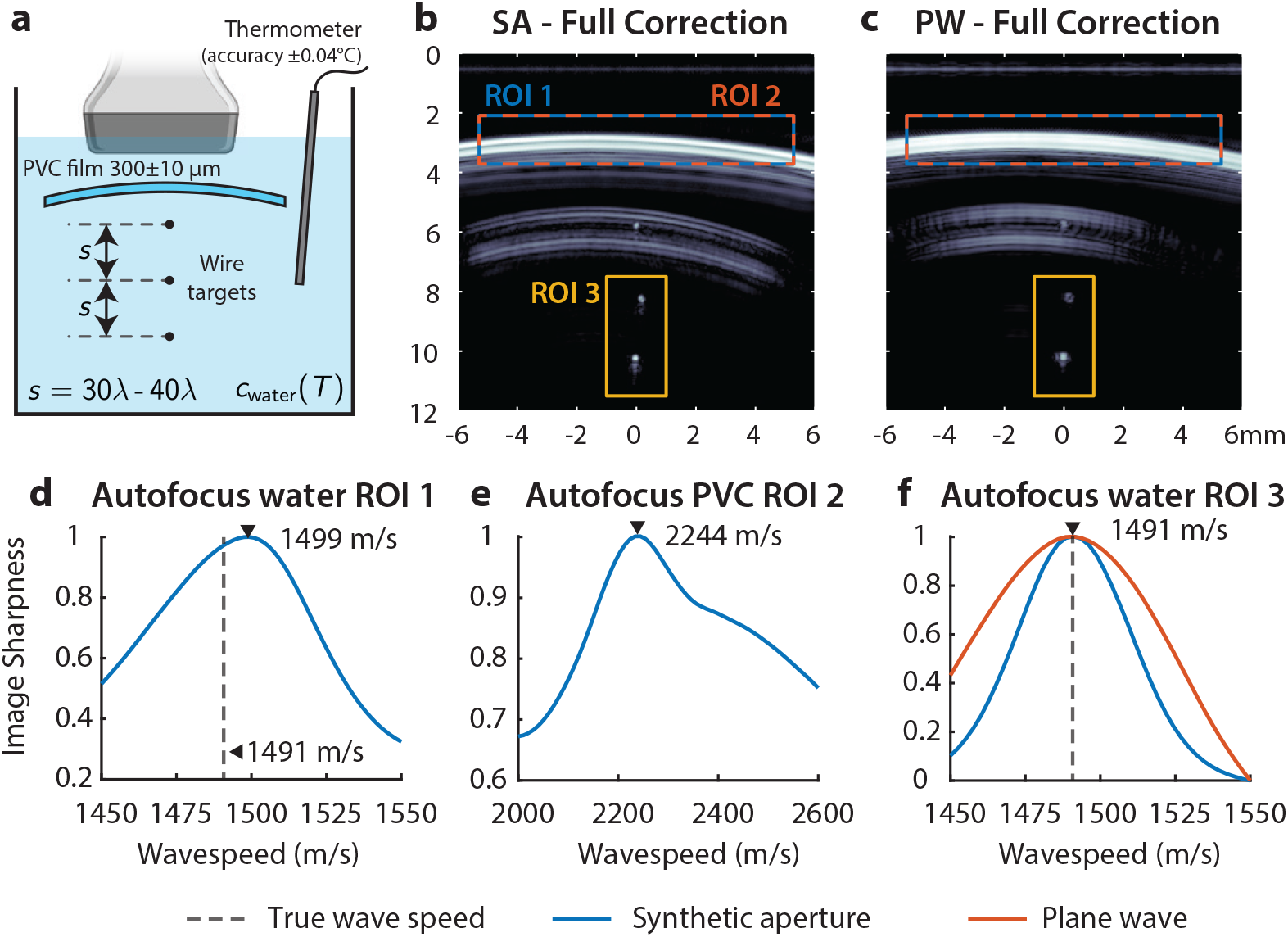
**(a)** Phantom validation setup. **(b)** The aberration-corrected synthetic aperture. **(c)** Aberration-corrected plane wave image of the plastic film and wires. **(d)** Autofocusing of the first water layer in ROI 1, using the synthetic aperture data. **(e)** Autofocusing of the PVC layer in ROI 2, using the synthetic aperture data. **(f)** Autofocusing of the last water layer in ROI 1, using both the synthetic aperture (blue line) and plane wave (orange line) data.

**Fig. 4.**
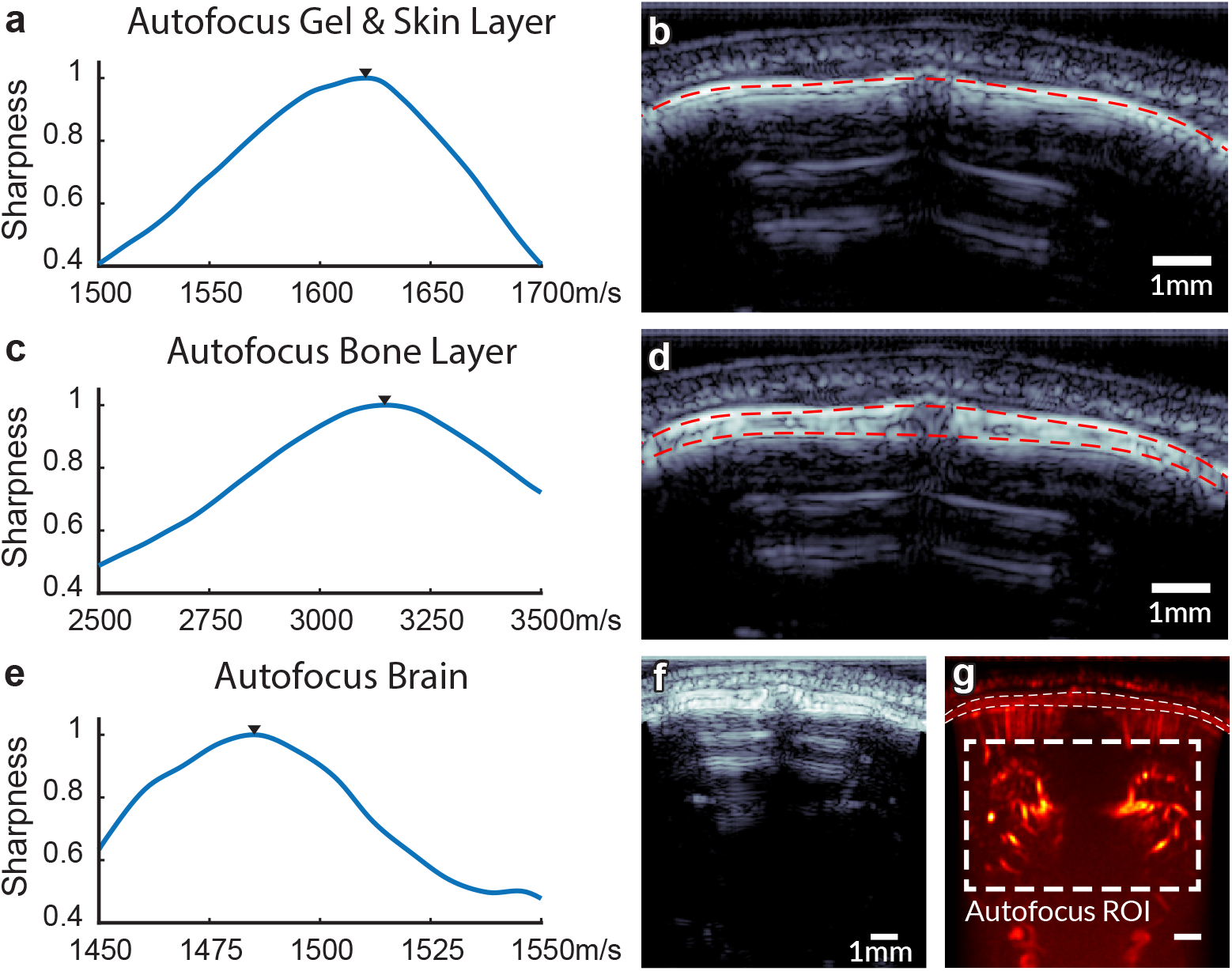
Wave speed estimation. **(a)** Result of autofocusing the gel and skin layer using the synthetic aperture data. The image is reconstructed using the 2-layer model. **(b)** The synthetic aperture image reconstructed with the estimated wave speed of the gel/skin layer. In this image, the outer skull surface is segmented using Dijkstra’s algorithm (red dashed line). **(c)** Result of autofocusing the skull bone layer using the synthetic aperture data and the 3-layer model. **(d)** The synthetic aperture image reconstructed with the estimated wave speeds. The inner skull surface is segmented, indicated by the lower red dashed line. **(e)** Finally, the brain angiography data is used with the 4-layer model to autofocus the wave speed in the brain tissue. **(f)** Aberration corrected plane wave B-mode. **(g)** Power Doppler data to estimate the brain wave speed.

Secondly, the wave speed in the skull bone is found using the 3-layer model. We define a ROI around the bone (Fig 4d) and maximize the sharpness of the inner skull surface. To avoid influence of the outer bone surface during maximization of the sharpness, a mask that mutes the outer bone surface is applied. After obtaining the wave speed of the skull bone, we can reconstruct a high resolution B-mode of the total skull, and segment the inner skull surface.

To segment the skull, we use Dijkstra’s algorithm to find the lateral path through the image that has maximizes brightness. This path is smoothed with a 3 sample moving average filter. Then, a piece wise cubic spline is fitted to the path. To have a precise segmentation, the image is reconstructed with an isotropic pixel size of 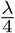.

To estimate the final missing velocity in the 4-layer model, we use the data from the brain angiography sequence. We reconstruct the brain using the 4-layer model and an initial wave speed of 1540 m*/*s, and form the Power Doppler image. In this image we define a ROI around the brain vasculature (Fig. 4g), and autofocus the ROI to maximize the sharpness of the Power Doppler images. We use the Power Doppler image to autofocus since the synthetic aperture image has too low SNR. Instead, we could have used the B-mode of the brain angiography sequence, but the presence of multiple reflections of the skull biases the autofocusing. Finally, we use the fully characterized 4-layer model to reconstruct the full aberration corrected brain angiographic image.

During the autofocusing process for brain tissue, the vascular content within the region of interest (ROI) changes depending on the tested wave speeds, which biases the focus quality metric. To mitigate this effect, the bounding box of the ROI was shifted by

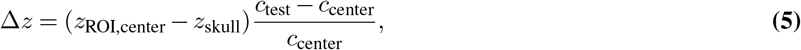

with *z*_ROI,center_ the depth of the center of the ROI for autofocusing, *z*_skull_ the depth of the inner skull surface at the center of the image, *c*_test_ the wave speed tested, and *c*_center_ the mean value of the tested wave speeds.

#### 2.4.6 Phantom validation

To validate our adaptive aberration correction technique, we estimated the wave speed of three distinct layers in a phantom setup. We submerged a wire phantom with 3 nylon wires with a diameter of 10 μm in a tank with degassed water (Fig. 3a). We positioned a 300 ± 10 μm PVC aberrating layer above the wires (measured with caliper; Mitutoyo, Kawasaki, Japan). The phantom was imaged using the skull segmentation and brain Doppler sequences. Before and after imaging, the water temperature was measured using a high accuracy thermometer with an accuracy of 0.04 °C (PT100 thermocouple, Greisinger electronics, Germany).

After imaging, we first autofocused the layer of water between transducer and PVC using the 2-layer model, then used the 3 layer model to find the wave speed in the PVC, and finally used the 4-layer model on the plane wave data to estimate the wave speed in the water around the wires. The temperature of the water was converted to the ground truth sound speed of the water using a calibration curve (62). When the ultrasound estimated thickness of the PVC layer and the wave speed of the water in the third layer match the ground truth, our method is validated.

### 2.5 Evaluation of image quality improvement

We use three reconstruction approaches to compare image quality. The reference image has no correction (NC) and relies on conventional DAS. We compare the reference images to a lens corrected image (LC) using the 2-layer model, and a full correction (FC) with the 4-layer model.

To evaluate the resolution improvement and the ability to resolve more vessels, we analyze the Doppler intensity profile along a line through the cortex. Since vessels rarely exhibit a prominence of at least 6 dB relative to adjacent local minima, the conventional full width at half maximum (FWHM) metric is not reliably applicable. Instead, we identify vessels with a minimum peak prominence of 2 dB, and compute their width at the -2 dB level.

To assess whether the improvement of our aberration correction technique varies over imaging depth, we compute the mean value and sharpness of the power Doppler image in three ROIs: cortex, midbrain and deep (Fig. 7a). We then determine the improvement with respect to the reference image, by computing the ratio for both the magnitude and sharpness of the power Doppler image, of LC and FC over NC. Since in the three reconstruction approaches the skull depth and apparent thickness is different, the ROIs are offset by the difference in apparent skull depth with respect to NC.

To further understand the effect of our aberration correction, we determined the phase aberration law for the different points in the image (see crosses in Fig. 7a). The aberration law was found by taking the difference between the return time of flight from the conventional DAS approach and the full correction.

Since the skull is reconstructed with too low a wave speed in the NC and LC images, the bone appears too thin. In addition, the bone layer acts as a diverging lens. When using our 4-layer model to correct for the aberrations, this leads to a lateral and axial shift of the brain vasculature in the FC image.

To quantify the thickness of the bone layer, we measured the distance between the outer and inner skull segmentations. We determined the minimum distance of the outer skull segmentation to the inner skull segmentation for every pixel to obtain the skull thickness as a function of lateral position. Finally, we compute the mean skull thickness per subject.

## 3 Results

In characterizing the transducer, we found a lens wave speed of 1009.6 m*/*s, a lens thickness of 518.5 μm, and a time to peak of 0.335 μs for the single cycle waveform and 0.467 μs for the three cycle waveform.

### 3.1 Phantom validation

In the validation experiment, the water temperature was 22.80 °C, corresponding to a wave speed of 1491 m*/*s (62). The results of autofocusing for each layer are shown in Fig. 3. The synthetic aperture imaging sequence was used to estimate wave speeds in the first water and PVC layers, resulting in 1499 m*/*s and 2244 m*/*s respectively. The segmentation of the PVC film revealed a thickness of 290 μm. Finally, autofocusing of the wire targets in the water layer below the PVC film yielded a wave speed of 1491 m*/*s for both synthetic aperture and plane wave imaging methods. The full aberration corrected phantom synthetic aperture image is shown in Fig. 3b, and the plane wave image in Fig. 3c. These findings validate our method’s ability to accurately estimate wave speeds and segment layer boundaries of a thin aberrating layer, under controlled experimental conditions.

### 3.2 Wave speed estimation

Fig. 4 shows the autofocusing curves and step by step B-mode images of the rat skull. Fig. 4a shows the image sharpness for the tested wave speeds in the gel/skin layer, where we found a mean wave speed of 1628±7 m*/*s for all rats. Fig. 4b shows the lens corrected synthetic aperture image of the rat skin and skull with the outer skull surface segmentation. In this image, the bone is reconstructed with a too low wave speed, and shows up with an underestimated thickness. Fig. 4c shows the bone autofocusing curve, resulting in a mean wave speed of 3247±110m*/*s for all rats. Fig. 4d shows the full aberration corrected synthetic aperture image of the rat skull, where the inner skull surface is segmented. Compared to the lens corrected image, the bone is now reconstructed with the correct wave speed and shows up with the correct thickness. The average thickness for all rats was 388±41 μm. Fig. 4e shows the autofocusing result for the brain, where we found a mean wave speed of 1526±55m*/*s for all rats. Finally, Fig. 4f,g shows the full aberration corrected plane wave B-mode of the rat skull and power Doppler of the brain vasculature. Due to the presence of multiple reflections in the B-mode, we used the shown autofocusing ROI in the Power Doppler to retrieve the brain wave speed. The results of the wave speed estimation and segmentation for each rat are shown in Table 1.

**Table 1.** Age, weight, the results of autofocusing for all layers, and the mean*±*SD segmented skull thickness for each rat.

| Rat | Age<br>(days) | Weight<br>(g) | Wave speed (m/s) | | | Thickness<br>skull ($\mu\text{m}$) |
| --- | --- | --- | --- | --- | --- | --- |
|  |  |  | Skin/Gel | Bone | Brain |  |
| 1 | 47 | 260 gr | 1626 | 3310 | 1590 | $352 \pm 74$ |
| 2 | 47 | 288 gr | 1622 | 3120 | 1486 | $378 \pm 77$ |
| 3 | 54 | 368 gr | 1636 | 3309 | 1503 | $433 \pm 99$ |

### 3.3 Improvement of image quality

Fig. 5 shows the Doppler images of the three rats, for no correction (NC), lens correction (LC) and full correction (FC). Vessels appear sharper and with higher intensity in FC compared to LC and NC, and blurry vessels are resolved after correction. In the FC images, there are dark areas on the sides of the images, which originate from the limited lateral extend of the segmentation.

**Fig. 5.**
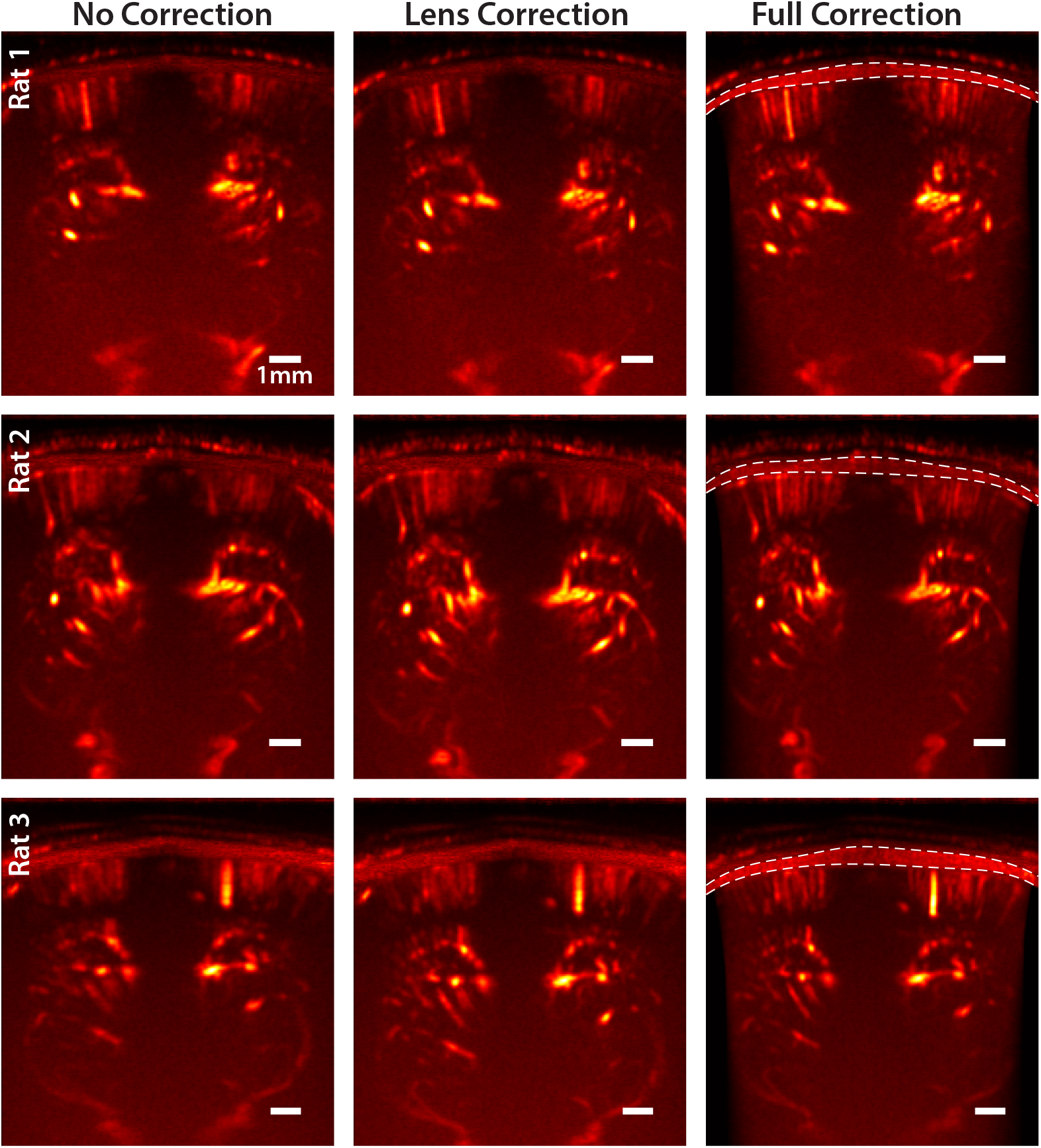
Power Doppler ultrasound images of three living rat brains. First column the uncorrected image using conventional DAS. The second column shows the lens aberration corrected image using the 2-layer model. The third column shows the fully corrected image using the 4-layer model. All scale bars denote 1 mm.

Fig. 6 shows a quantitative analysis of the resolution improvement in the cortex due to aberration correction. Fig. 6d shows that the full correction allows to individually resolve more vessels in the cortex, compared to no correction and lens correction. This finding was consistent for all three rats. In addition to resolving more vessels, Fig. 6e shows that vessel width (width at -2dB level) decreases in LC and FC, indicating an improvement in lateral resolution. For rat 1 the median vessel width was 239.7 μm for NC, 213.2 μm for LC and 155.7 μm for FC. For rat 2 the median vessel width was 217.5 μm for NC, 224.0 μm for LC and 162.6 μm for FC. For rat 3 the median vessel width was 256.5 μm for NC, 208.7 μm for LC and 149.7 μm for FC. On average, the improvement in vessel width was 9% for LC and 32% for FC.

**Fig. 6.**
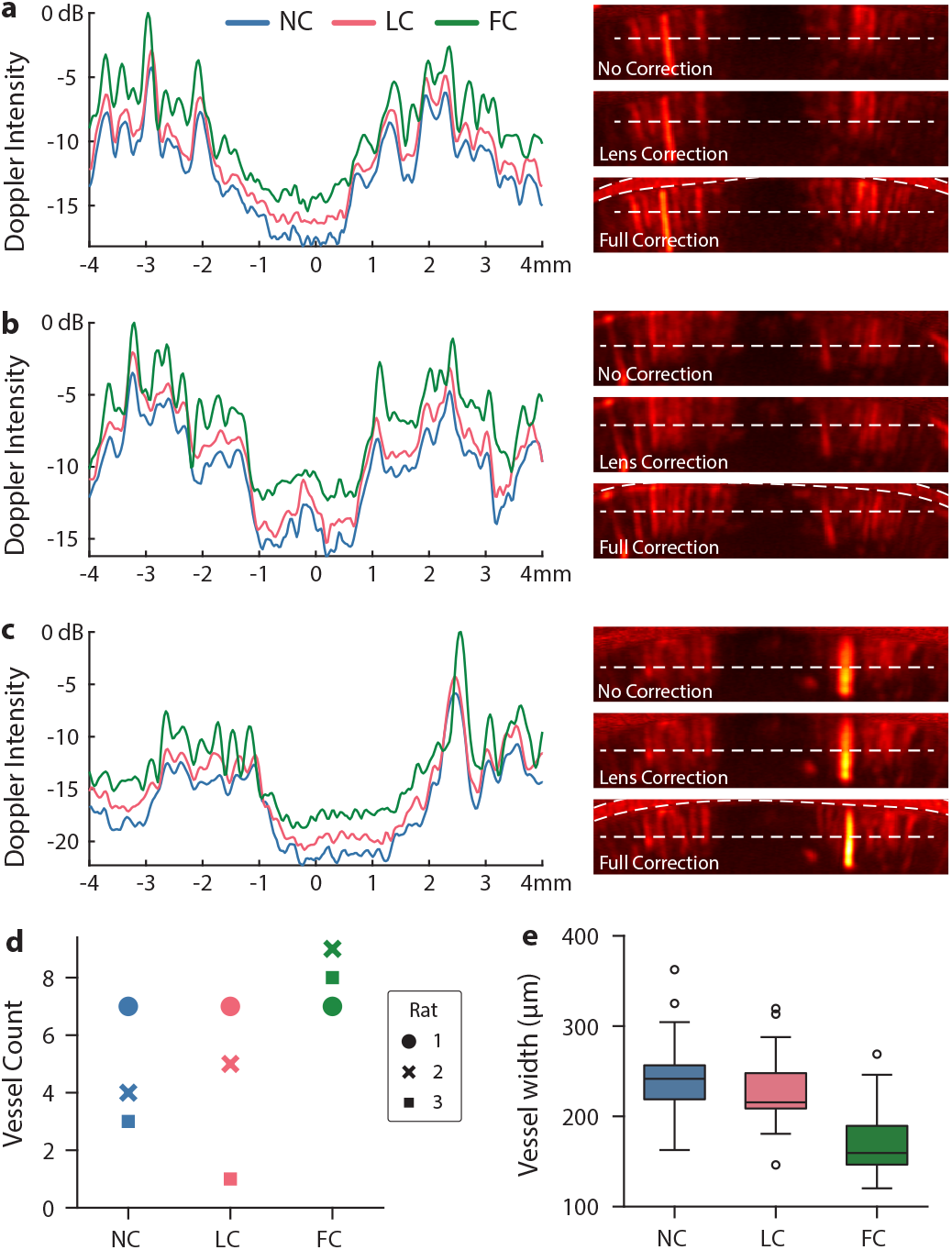
Power Doppler profiles through the cortex, of rat 1. **(a)**, rat 2 **(b)** and rat 3 **(c)**. Colors indicate the different reconstruction approaches, with in blue No Correction (NC), in red Lens Correction (LC) and in green Full Correction (FC). The profile corresponds to the white dashed line overlaid on the cortex zooms on right of the profiles. Panel **(d)** shows a vessel count for the three reconstruction approaches. The vessel count is obtained by identifying peaks in the Doppler intensity profile with a minimum prominence of 2 dB relative to adjacent local minima. **(e)** Shows the vessel width, defined as the width at the -2 dB level for peaks found in the power Doppler image. The box plots show the combined data of the three rats.

Fig. 7 shows the improvement of the aberration correction method as a function of imaging depth. Fig. 7b shows the aberration law for three different depths in the image, corresponding to the cortex, midbrain and deep brain and both brain hemispheres. It becomes clear that the aberration is the strongest for the ROI closest to the aberrating layer, and decreases with imaging depth. The maximum time difference of the aberration law is≈ 1.15 μs, which corresponds to approximately 16 ultrasound periods at 13.9 MHz. Fig. 7c,d show the relative improvement in magnitude and sharpness of the power Doppler image for the three ROIs at different depths. The relative magnitude improvement for LC was 1.3 in the cortex, 1.5 in the midbrain and 1.4 in the deep brain. For FC the relative magnitude improvement was 2.1 in the cortex, 1.8 in the midbrain and 1.6 in the deep brain. The relative sharpness improvement for LC was 3.9 in the cortex, 8.2 in the midbrain and 4.5 in the deep brain. For FC the relative sharpness improvement was 31.8 in the cortex, 19.8 in the midbrain and 8.4 in the deep brain. The improvement in magnitude and sharpness of the FC power Doppler image is highest for the cortex, and decreases with imaging depth. The improvement in FC is higher than in LC for all ROIs and depths.

**Fig. 7.**
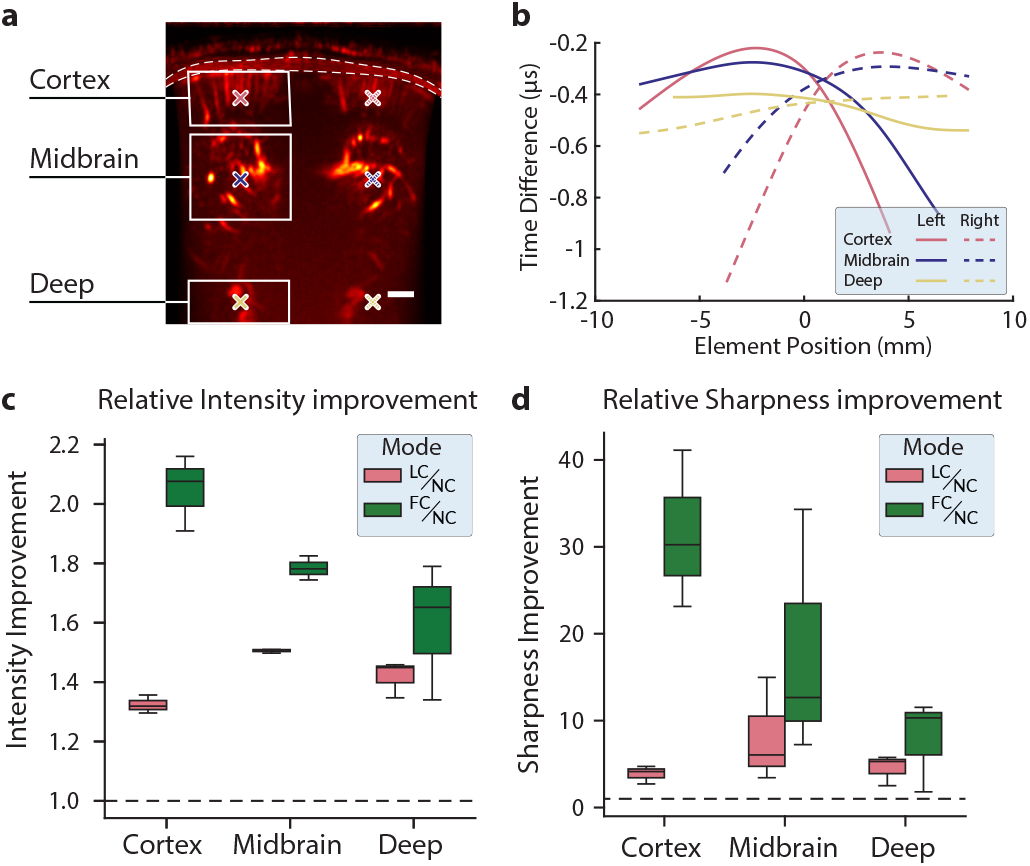
Aberration correction performance across depth. **(a)** Three regions of interest (ROIs) at different depths in the image, corresponding to the cortex, the midbrain and deep. The scale bar denotes 1 mm. **(b)** Aberration law for three different depth (colors correspond to the markers colors in **(a)**). Solid lines denote the left hemisphere and the dashed lines the right hemisphere. The aberration law is found by the different in return time of flight between the No Correction and Full Correction travel time lookup tables. **(c)** Relative improvement in magnitude of the Lens Correction (LC) and Full Correction (FC) power Doppler images, with respect to No Correction (NC) for the three ROIs at different depth. The black dashed line at 1.0 indicates no improvement. **(d)** Relative improvement in the Doppler sharpness for three ROIs at different depth of Lens Correction (LC) and Full Correction (FC) with respect to No Correction (NC).

## 4 Discussion

### 4.1 Estimation of tissue wave speed and skull thickness

In this study, we report an adaptive 4-layer ray-tracing method approach to correct skull aberrations in transcranial ultrasound imaging of the adult rat brain. We first validated on a phantom experiment with an aberrating layer having known thickness and a known wave speed in water. The wave speed in water was estimated with an accuracy equal or better than 0.56%. The wave speed found for the PVC layer is in line with the reported values in the literature (63), and the estimated thickness retrieved by segmenting the PVC layer in the B-mode is within the error margin of the caliper measurement (*±*10 μm).

The estimated wave speed in the gel/skin layer and the bone layer was in agreement with the literature (50, 52). The variability in wave speed may originate from subject variability in age, diet, or experimental conditions such as the probe-skull angle and the fact that the inner and outer skull surface may not be perfectly parallel, reducing specular reflection. The skull thickness retrieved from the aberration corrected synthetic aperture images matched well with earlier reports on body mass and skull thickness in rats (54). For the brain, the estimated wave speeds showed larger standard deviation, but were still in line with expected values from the literature.

### 4.2 Depth-dependence of aberration and its correction

Evaluating the impact of aberration correction as function of imaging depth, showed that the correction performed best in the cortex, as evidenced by magnitude profiles of the power Doppler image through the cortex (Fig. 6), along with the magnitude and sharpness ratios of the power Doppler images (Fig. 7c,d). For all imaging depths, both the fully corrected (FC) and lens corrected (LC) methods outperformed the non-corrected (NC) approach. As imaging depth increased, the performance of FC became closer to that of LC. This trend can be explained by the nature of aberrations, which are the strongest near the aberrating layer where the incident angles are steepest. This observation aligns with the aberration law shown in Fig. 7b, which demonstrates that as depth increases, the aberration law flattens. The flattening explains why the performance difference between LC and FC diminishes with depth. Closer to the aberrating layer, the aberration law also reveals that it is not possible during ray-tracing to find a path to all receiving elements (indicated by pink lines in Fig. 7). This is caused by steep incident angles at the bone interface, which lead to critical reflection, as shown in Fig. 1b. For deeper regions, the incident angles at the bone interface decrease, and the ray-tracer successfully can find a path for all receiving elements.

### 4.3 Challenges in skull segmentation and alignment between transducer and skull

The main challenge of the aberration correction method introduced in this manuscript lies in skull segmentation. For young animals with very thin skulls, the axial resolution of the imaging system can become insufficient and prevent the accurate delineation the outer and inner skull interfaces. For example, our validation experiment demonstrated that a 300 μm aberrating layer could be resolved using a 1-cycle imaging pulse at 15 MHz, but the 3-cycle long plane wave Doppler sequence had an insufficient resolution to detect outer and inner interfaces of the PVC layer. This highlights the need for a high resolution sequence for skull segmentation. In the current implementation, the axial resolution of our imaging system allows imaging of rats weighing 200 g or more approximately (54).

The accuracy of the skull segmentation also depends on obtaining clear specular reflections from both the outer and inner surfaces of the bone. The outer specular reflection could be reliably obtained during the *in vivo* data acquisition by verifying with live imaging. However, achieving a clear specular reflection from the inner surface proved challenging due to the absence of an aberration-corrected preview. Obtaining a specular reflection from the inner skull surface is even more challenging when the skull surfaces in the imaging plane are not parallel.

Implementing the method in 3D using a 2D matrix array would offer significant advantages. In 3D imaging, the requirement for precise probe positioning to acquire specular reflections is virtually eliminated. Furthermore, 3D imaging allows imaging of the brain below non-parallel skull surfaces.

### 4.4 Implications for transcranial functional imaging

The advancements presented in this study have significant implications for 2D fUS. A remaining challenge in 2D transcranial fUS is navigating to brain regions of interest. The requirement of positioning the probe perpendicular to the skull, often results in a power Doppler image of the brain that differs significantly from the standard brain atlas.

One key advantage of our method is the ability to retrieve the wave speed model once and apply it across any other acquisition sequence. After initialization, functional recordings can be aberration corrected at no additional computational cost, provided that the transducer stays in place.

Our method improves the reconstruction accuracy to show brain regions at the correct anatomical locations and with the correct shape, thus facilitating accurate comparisons with anatomical and functional data from other imaging modalities. Furthermore, enhanced spatial resolution in cortical regions substantially improves the spatial specificity of the fUS signal, improving the usability of transcranial fUS in functional connectomics (64).

Our improved resolution in the cortex would make fUS more sensitive to weak and or transient neural activity. For instance, to create accurate somatotopic mappings during whisker stimulation experiments. The barrelfield is located in the primary somatosensory cortex, and contains distinct neuronal clusters, known as barrels, which correspond one-to-one with the whiskers on the rat’s face. With inter-barrel distances of approximately 50 − 100 μm, our enhanced resolution would allow the detection of single-whisker activations (65). Similarly, our method could achieve higher resolution retinotopic mappings of the visual system (66) and tonotopic mappings of the auditory system (67).

To fully unlock the potential of noninvasive fUS and overcome the current limitations of this study, future efforts will focus on extending our aberration correction method to 3D imaging. This advancement would enable 4D transcranial imaging, offering comprehensive spatio-temporal resolution and further advancing the utility of fUS in neuroscience.

## 5 Conclusion

We developed an adaptive aberration correction method using 4-layer ray-tracing that can improve image quality and signal spatial specificity in transcranial ultrasound Doppler imaging of the adult rat brain. The method adapts to each subject and each brain section being imaged without requiring prior information from additional imaging modalities like a CT or MRI scan. The method consistently outperformed conventional delay-and-sum image reconstruction, and holds promise for functional imaging applications that require high spatial sensitivity.

## 6 Acknowledgment

This project was partially funded through grants from the Medical Delta Ultra HB program (RW), the Chan Zuckerberg Initiative (Dynamic RFA number 2023-321233, EMI), European Union’s Horizon 2020 research and innovation programme grant, ERC-StG ‘HelpUS’ 758703 (FN, VG), the European Union (Marie-Sklodowska Curie Fellowship MIC-101032769, BH) and the 4TU Precision Medicine Program (DM).

We thank Ruud van Tol (NIN, KNAW) for designing the ultrasound probe holder, Henry den Bok (ImPhys, TU Delft) for fabricating wire phantoms, Ronald Ligteringen (ImPhys, TU Delft) for facilitating access to computational resources, and Gert Jan de Fluiter (NIN, KNAW) for his assistance with animal care.

